# Neural decoding dissociates perceptual grouping between proximity and similarity in visual perception

**DOI:** 10.1101/2021.11.15.468580

**Authors:** Lin Hua, Fei Gao, Chantat Leong, Zhen Yuan

## Abstract

Previous research on perceptual grouping primarily focused on the dynamics of single grouping principle in light of the Gestalt psychology. Yet, there has been comparatively little emphasis on the dissociation across two or more grouping principles. To tackle this issue, the current study aims at investigating *how*, *when*, and *where* the processing of two grouping principles (proximity and similarity) are established in the human brain by using a dimotif lattice paradigm and adjusting the strength of one grouping principle. Specifically, we measured the modulated strength of the other grouping principle, thus forming six visual stimuli. The current psychophysical results showed that similarity grouping effect was enhanced with reduced proximity effect when the grouping cues of proximity and similarity were presented simultaneously. Meanwhile, electrophysiological (EEG) response patterns were able to decode the specific pattern out of the six visual stimuli involving both principles in each trail by using time-resolved multivariate pattern analysis (MVPA). The onsets of the dissociation between the two grouping principles coincided within three time windows: the earliest proximity-defined local visual element arrangement in the middle occipital cortex, the middle-stage processing for feature selection modulating low-level visual cortex in the inferior occipital cortex and fusiform cortex, and the higher-level cognitive integration to make decisions for specific grouping preference in the parietal areas. In addition, brain responses were highly correlated with behavioral grouping. The results therefore provide direct evidence for a link between human perceptual space of grouping decision-making and neural space of these brain response patterns.

**Significance statement:** How does our brain aggregate meaningless local elements to meaningful global patterns? Here, we used visual stimuli that can be perceived as a whole orientation from one of the two grouping cues (proximity and similarity) when they were presented simultaneously. Behavioral responses showed a trade-off between proximity and similarity. Further, time-resolved MVPA and source estimation results showed three stages for the processing of two grouping cues, involving an early stage in the low-level visual cortex, then the middle stage in the lateral visual cortex, and finally the late stage in the parietal areas. This was among the first attempt employing multivariate analysis to decode the dissociative processing of both proximity and similarity principles within one single study.

## Introduction

The perceptual grouping or organization depicts the process when the human brain aggregates meaningless local sensory elements to meaningful global patterns. Gestalt psychologists suggested that perceptual grouping is based on the mechanism sensitive to various principles, such as proximity, similarity, good continuation, closure, and common fate of visually ordered elements in our real-world experience (<u>Wertheimer, 1922</u>; <u>1923</u>). Among those principles, proximity and similarity are the earliest discovered phenomena and also recognized as the most fundamental processes (<u>Wagemans et al., 2012</u>). Previous research has established the dynamic signatures and anatomical distributions for single grouping principle in vision science. Compared to ungrouped local elements in the space, single perceptual grouping principle implicates an early visual processing (<u>Nikolaev et al., 2008</u>; <u>Kurylo et al., 2017</u>). Meanwhile, widespread brain activation was identified, including areas V1 (<u>Wannig et al., 2011</u>; <u>Stoll et al., 2017</u>), V2 (<u>Merigan et al., 1993</u>; <u>Murray et al., 2004</u>), the lateral occipital complex (LOC; <u>Fang et al. (2008)</u>) of the visual cortex, the middle temporal cortex, the inferior parietal cortex, and the prefrontal cortex (<u>Seymour et al., 2008</u>; <u>Carther-Krone et al., 2020</u>).

However, while human beings usually face the simultaneous processing of two or more principles of perceptual grouping information in natural environment, what is not yet clear is its underlying mechanisms. Existing behavioral studies observed that grouping effects were enhanced by the cooperation of two grouping principles, while the effects decreased when the two principles were competing (<u>Quinlan and Wilton, 1998</u>; <u>Luna and Montoro, 2011</u>). Yet, those findings were mostly based on the descriptions of conscious experience regarding the units that people naturally perceive, instead of manipulating these grouping principles in fine-grained psychophysical settings. This limitation leads to a failure in quantifying the effects of two or more grouping principles and detecting the specific principle in operation.

To identify the neural correlates of differing grouping principles, extant imaging studies performed univariate analysis [e.g., event-related potential (ERP) analysis] to differentiate the two grouping principles generated by separate stimuli. Some studies (<u>Han, 2004</u>; <u>2005b</u>) suggested that grouping by proximity, relative to shape similarity, is consistently associated with enhanced positive ERP amplitudes peaking around 100 ms after stimulus onset over occipital electrodes (P100) and around 300 ms in central and parietal electrodes (P300). Additionally, grouping by similarity relative to proximity elicited increased amplitudes in a temporo-occipital negative component (N200). In contrast, other studies (<u>Luna et al., 2016</u>; <u>Villalba-García et al., 2018</u>) reported a null P100 effect regarding the difference between proximity and similarity grouping. This discrepancy might be attributed to the limitation of phenomenological paradigm and univariate analysis approach. In the aforementioned ERP studies, only components containing significant difference between proximity and similarity grouping were measured, while those were discarded whose signals could actually distinguish different perceptual grouping states yet failed to reach a significance (<u>Cichy and Pantazis, 2017</u>).

To bridge the theoretical and methodological gap, the current study aims at directly quantifying the relationship of two basic grouping principles (proximity and similarity) presenting simultaneously by using a dimotif lattice paradigm. Specifically, both psychophysical and spontaneous electrophysiological (EEG) data were recorded to measure the precise value of the different perception between proximity and similarity grouping. To resolve the limitation of EEG univariate analysis, we innovatively performed time-resolved multivariate pattern analysis (MVPA) to dissociate two grouping principles. This approach would include the interaction between multiple channels/trails, so as to detect the subtle changes in brain activation patterns associated with proximity and similarity grouping in the experiment. Additionally, source estimation was calculated to examine how the neural mechanisms of two grouping principles were integrated in the spatial domain. We hypothesized that grouping effects of two principles would be precisely quantified when manipulating the strength of one grouping effect and controlling the other. Further, the dissociative processing of two grouping principles would be verified in three time windows, implicating differing stages in both temporal and spatial domains.

## Materials and Methods

### Participants

Thirty college students (19 males, mean age: 21.4 ± 2.8 years) from the University of Macau participated in the experiment and received financial reward. All participants reported no neurological illness or mental disorder, and were right-handed with normal or corrected-to-normal vision. Informed written consent was obtained from each participant prior to the experiment. Four participants (3 males and 1 female) were excluded from the formal analysis due to not following the task instructions (pressing the same button throughout the whole test) or more than 10% EEG artifacts. The research was approved by the Institutional Review Board at the University of Macau.

### Dimotif lattice stimuli

Two categories of motifs (i.e., squares and discs, <u>Kubovy and Van Den Berg, 2008</u>) were used as visual stimuli in the dimotif lattice (<u>Grünbaum and Shephard, 1987</u>), which could help quantify the interaction between proximity and similarity grouping effects. The arrangement of all motifs was constrained by three parameters (*a*, *b*, and *γ*), as demonstrated in Figure 1A. Specifically, *a* and *b* denote the distances of neighboring motifs along the two main axes, which are perpendicular to each other. All elements in either string of direction *a* consist of the identical motifs, while adjacent strings are made of heterogeneous motifs. Yet, each element string in direction *b* consists of alternating occurrences of the two motifs, whose pattern remains the same across all strings. The last parameter *γ* represents the angle between *b* axis and the horizontal line (measured counterclockwise).

**Figure 1.**
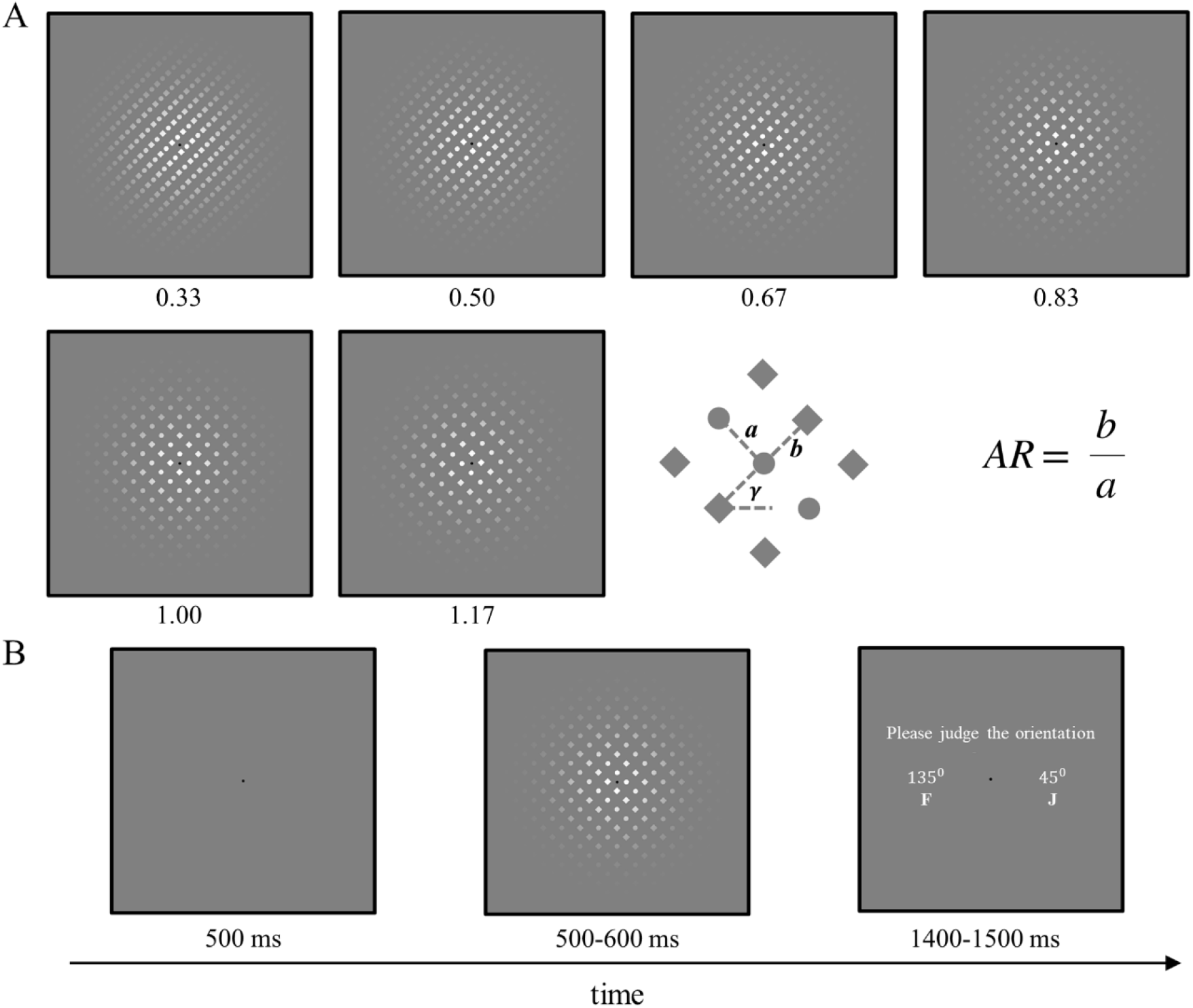
Experimental design and procedure. (A) Visual stimuli consist of six types of dimotif lattice pattern with a coherent (45° or 135°) or ambiguous global orientation, as a result of altered AR values (0.33, 0.50, 0.67, 0.83, 1.00 and 1.17). The calculation formula and diagram of the AR are presented in the lower right. (B) Timing and sequence of stimuli in an experimental trail. Participants were instructed to press “F” key by the left index finger for a 135° global perceived orientation and to press “J” key by the right index finger for a 45° perception.

The orientation of the dimotif lattice patterns could therefore be defined by the aspect ratio 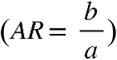 and *γ*, given a fixed *a* value. In the current design, *a* and *γ* were set to 1.5° and 45°, respectively, while AR was chosen from 0.33, 0.50, 0.67, 0.83, 1.00 and 1.17, thus forming six dimotif lattice patterns. To evaluate participants’ grouping preference, the perceived orientation of 45° would be seen as a proximity, while 135° as the similarity grouping preference.

### Procedure

Participants were seated 55 cm from the display with a maximum visual angle of 25 × 25 degrees. Visual stimuli were generated by a Dell 64 bit-based machine (12G RAM) with an AMD Radeon HD graphics card running Psychtoolbox-3 software (www.psychtoolbox.org) on Windows Professional 7. All motifs (squares and discs) were presented with a diameter of 1° against the gray background on a Dell P2312H monitor with a resolution of 1024 × 768. The element-background contrast was 150% (Weber contrast, *c* =(*I* − *I*_*b*_)/*I*_*b*_, where *I* and *I*_*b*_ were the luminance of the motifs and the background respectively). The whole pattern was masked with a Gaussian mask (20° diameter), at whose center was a black point as fixation (0.5° diameter). The point fixation stayed on the screen for the whole test and participants were asked to fixate at it. Within each trial, a blank with the point fixation was presented for 500 ms, followed by a dimotif lattice pattern lasting for 500 ms with a jitter of 100 ms. After the dimotif lattice disappeared, participants would encounter a probe asking them to judge the orientation of the previous pattern within 1400-1500 ms (Figure 1B). Participants were instructed to perform a two-alternative forced choice (2AFC) for two orientations (45°or 135°) by pressing the corresponding buttons labeled in the keyboard. Each of the six dimotif lattice patterns was presented for 80 times, thus making 480 trials in a random order. Each participant completed three blocks with 160 trials in each block.

### Psychophysical analysis

Psychophysical performance was quantified as the percentage of dimotif lattice pattern with a response of 135° (i.e., similarity grouping preference). Psychometric functions were generated by fitting a cumulative Gaussian sigmoid curve by using the Psignifit toolbox (version 4, https://github.com/wichmann-lab/psignifit/wiki), which implements the maximum-likelihood method for parameter estimation, and finds confidence intervals by bias-corrected and accelerated bootstrap method (<u>Schütt et al., 2016</u>). The point of subjective equality (PSE) for each participant was extracted from the horizontal axis value corresponding to 0.5 on the vertical axis of the psychometric curve. The statistical visualization of PSEs across participants were performed by using the boxplot function in MATLAB (Mathworks, Natick, MA, USA).

### EEG recording and preprocessing

Participants were seated in a sound-attenuated and electrostatically shielded room during the experiment. The continuous EEG data were recorded from a 64-channel Biosemi Active Two EEG amplifier system (Biosemi, Amsterdam, Netherlands) with Ag/AgCl scalp electrodes placed according to the international 10-20 system on an elastic cap. During the online acquisition, EEG data were sampled at 2048 Hz with a bandpass filter of 0.01-200 Hz. The input impedance of all channels was kept below 5 kΩ.

The EEG data were processed offline by using custom-made scripts in MATLAB, the EEGLAB toolbox (<u>Delorme and Makeig, 2004</u>), and sLORETA software (<u>Pascual-Marqui, 2002</u>) for source analyses. After down-sampling the data to 500 Hz, a built-in fourth-order Butterworth band-pass filter was applied with cutoff frequencies between 0.15 Hz and 40 Hz. Then, epochs lasting from 100 ms before the stimuli onset to 500 ms afterwards were extracted, among which those with unique, non-stereotypic artifacts were discarded. Independent component analysis (ICA) was then performed, and components representing common ocular or cardiac artifacts were visually identified and removed for further analysis. Overall, less than 10% of all trials were rejected. Finally, data were re-referenced to the grand average of whole head.

### Global electric field analysis

Modulations in the strength of the electric field at the scalp were assessed using the global field power [GFP; <u>Murray et al. (2008)</u>]. GFP is calculated as the square root of the averaged squared voltage value recorded at each electrode, which could index the spatial standard deviation of the electric field at the scalp. Larger GFP value denotes stronger electric field. Differences in GFP waveform data was analyzed as a function of time when the data post stimulus onset was significantly different from the baseline among six dimotif lattice conditions. GFP amplitude was considered significant if it exceeded a 95% confidence interval for at least 20 ms consecutively relative to the baseline of 100-ms pre stimulus. Subsequently, GFP peaks were determined from the averaged GFP waveform across participants.

Topographic modulations were identified using randomization statistics applied to global map dissimilarity measures [GMD; <u>Murray et al. (2008)</u>]. GMD is calculated as the root mean square of the difference between strength-normalized vectors. Difference in GMD values was also analyzed as a function of time, using stimulus type as a within-subject factor. The statistics of GMD were validated by a topographic ANOVA with 5000 permutations (*p* < 0.05). Notably, GMD is independent of field strength and a significant GMD is indicative for different neural generators across six dimotif lattice conditions. Significant results of GMD were corrected by a duration threshold (20 ms).

### Source estimation

Global electric field analysis showed that the dissociative information of two perceptual grouping was available in three time windows and the GFP peaks at 108 ms, 236 ms and 310 ms corresponded to the latency of the P1/N1, P2 and N3 in the visual stimuli-evoked potential. The neural sources of the P1/N1, P2 and N3 were reconstructed for the significant period of GMD around each latency by using the sLORETA software, which provides current density values of 6239 voxels (5×5×5 mm resolution), modeled in the CIT168 brain atlas (<u>Pauli et al., 2018</u>). This method is a Laplacian-weighted minimum norm algorithm with no priori assumption about a predefined number of activated brain regions, thus constituting a more open solution to the EEG inverse problem.

The group differences at each component were examined by voxel-by-voxel single *t*-test statistics. T-test thresholds were computed using the built-in program of the sLORETA software. Correction for multiple comparisons were performed using a randomization test of statistical non-parametric mapping (SnPM) with 5000 randomizations.

### Multivariate pattern analysis

MVPA reveals topographic weightings of EEG signals that maximally distinguish perceptual grouping states within a given time interval. Here, a linear classifier was used based on L2-regularized logistic regression (<u>Fan et al., 2008</u>) to find the optimal projections of the sensor space for discriminating among six dimotif lattice patterns (one-vs-all decoding) or between two dimotif lattice patterns (one-vs-one decoding) at a specific time point (Figure 5A). This allowed the assessment of how and when the dissociation of perceptual information between proximity and similarity was available in the stimulus-locked EEG data. The timing of this availability was evaluated by using time-resolved decoding, in light of previous visual MEG/EEG studies (<u>Crouzet et al., 2015</u>; <u>Cauchoix et al., 2016</u>).

The accuracy of the classifier (linear L2-regularized logistic regression) fed with the multi-electrode single-trail EEG signal was evaluated for each time point independently. For each time point, the performance of the classifier was determined by using a Monte-Carlo cross-validation (CV) procedure (n=100), in which the entire data was randomly partitioned into 10 portions including a training set (90% of the trails) and a test set (the remaining 10%). Here, the cost parameter C used the default value of 1 for all analysis. These time windows were centered on and shifted from −100 to 500 ms relative to stimulus onset on stimulus-locked data.

Using the procedure above, a temporal generalization matrix [TGM; <u>Carlson et al. (2013)</u>; <u>Cichy et al. (2014)</u>]was obtained using one-vs-one decoding for each dimotif lattice pattern at each time point. Each TGM corresponded to a set of weights allowing optimal discrimination for each group at a given time point. By training at one time point and testing across other time points, the results could assess how each TGM represented different time points. Significance of a time-point was assessed using a paired *t*-test of the distribution (across participants) of decoding performance obtained using true versus shuffled labels. The results were corrected for multiple comparisons using a false discover rate (FDR) and a significance level of *p* = 0.05.

For each participant, decoding accuracy was approximated according to the averaged performance across CVs. Error bars in the analysis correspond to the non-parametric 95% confidence intervals of the mean obtained via bootstrapping. For each time point and CV, a measure of chance performance was obtained by performing an identical classification analysis using randomly permuted labels. At the single-subject level, classification accuracy was considered above chance when it was higher than classification accuracy obtained from permuted labels (paired *t*-test, α=0.05). The group analysis was performed following the same procedure, except that the group averages were computed across single-subject averages. Correction for multiple comparisons were used for a time-cluster-based approach, in which a time point was considered significant only when it was a member of a cluster of at least ten consecutively significant time points (i.e., 20 ms).

### Representational similarity analysis

Representational similarity analysis [RSA; <u>Kriegeskorte et al. (2008)</u>] was based on the multi-class decoding analysis results. For multi-class decoding, classifiers were trained to discriminate among six-ARs of dimotif lattice patterns, whose results were similar to the participants’ psychophysical performance. This analysis constructed a dynamic neural representational similarity matrix (RSM) for each subject and time point. For each time point, there was a 6×6 confusion matrix, where each cell represents the proportion of trials. Given one cell from the matrix, dimotif lattice pattern X was presented and the classifier categorized the trial as dimotif lattice pattern Y. The confusion matrix was then transformed into RSM. These matrices denoted representation of the neural space in which dimotif lattice pattern categories were coded, since they could show which dimotif lattice pattern-evoked neural patterns were similar. To construct the behavioral RSM, the percentage results of dimotif lattice stimuli were compared to each other and converted to Euclidean distance metric. Then, the representational similarity between dimotif lattice patterns could be compared by calculating Pearson correlations. The behavioral RSM was symmetrical along its diagonal, and the off-diagonal areas indicated the representational similarity across six dimotif lattice patterns. In particular, this behavioral RSM could estimate the participants’ perceptual space to proximity or similarity grouping preference. Then, the Pearson correlation coefficients were computed for each participant and each time point separately across the 36 cells of the neural and behavioral RSM. This correlation would therefore provide an estimation of the representational similarity of perceptual grouping in perceptual space and neural space.

## Results

### Psychophysical results

Psychophysical performance was measured as the percentage of trails perceived as 135° (i.e., similarity preference). The averaged percentage of behavioral responses across all participants for each dimotif lattice pattern (ARs from 0.33 to 1.17) were 7.62 ± 1.28%, 32.52 ± 2.67%, 53.75 ± 3.36%, 83.07 ± 2.12%, 91.97 ± 1.69%, and 95.88 ± 1.01%, respectively. Fitting the behavioral data using psychometric functions, it was found that as the AR increases, the percentage of similarity preference shows an S-curve growth tendency (Figure 2A). This suggests that participants were inclined to group discrete motifs into parallel strings in the direction *a* when the distance between two motifs increased in the direction *b*.

**Figure 2.**
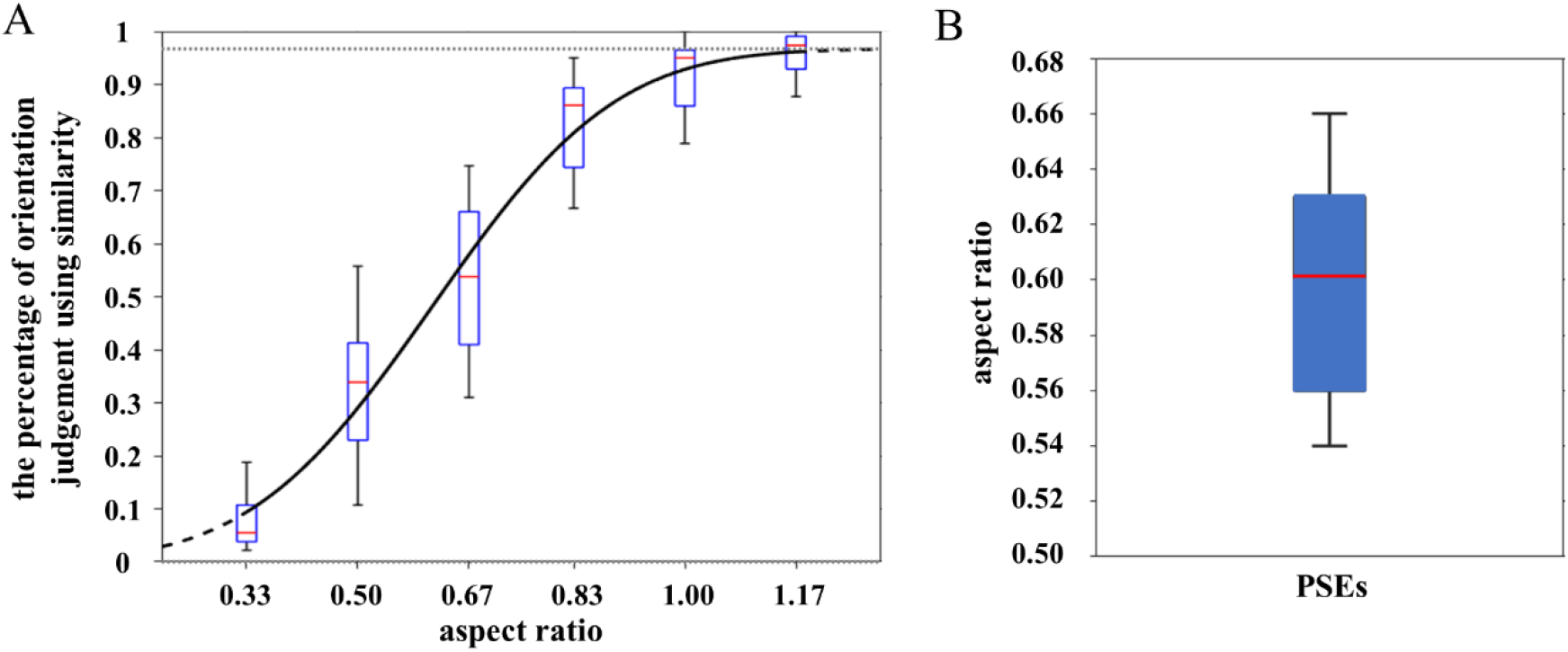
The fitted psychometric functions and averaged PSEs of all participants. (A) Illustrations of behavioral response between proximity and similarity grouping preferences among six dimotif lattice patterns. The cumulative Gaussian sigmoid curve was used to the fitted behavioral data with 95% confidence intervals of individual participant (dotted line). The S-curve was averaged across all participants. (B) Boxplot with whiskers representing PSEs. The box extends from the 25^th^ to 75^th^ percentiles while the whiskers extend from minimum to maximum values. The horizontal red line indicates the mean of PSEs (n=26). The PSEs indicate that the AR for participants has the same global perceived orientation for proximity and similarity grouping.

The experimentally obtained PSEs were averaged across all participants, yielding an AR of 0.601 ± 0.011 (Figure 2B). This result suggested that subjectively equated proximity effect of two perceptual grouping stimuli does not guarantee similar effects in the visual system due to the modulation of similarity effect. Converting PSEs to visual angle and computing the effect gap between proximity and similarity grouping by formula 1, similarity approximates 1.60° proximity in the current study.

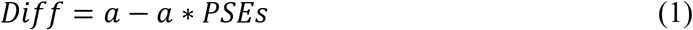

### Global electric field results

The neural dissociative processing of grouping preference between proximity and similarity evoked a significant electrophysiological response starting from 84 ms (GFP) relative to the pre-stimulus period (Figure 3A). Accordingly, the GFP showed the earliest signal increase for each dimotif lattice pattern (AR from 0.33 to 1.17) were 84 ms, 90 ms, 84 ms, 86 ms, 90 ms, and 84 ms. Three GFP peaks were identified among six dimotif lattice conditions at 108 ms, 235 ms, and 310 ms, corresponding to the visual stimulus-evoked potentials P1/N1, P2, and N3, respectively. The significant differences in the topographical distribution of the electric field independent of the electric field strength (GMD; *p*s < 0.05 lasting for 20 ms) revealed that the underlying neural generators varied between dimotif lattice stimuli at the following three time windows. The first stage started from 86 ms to 130 ms after stimulus onset with a negative potential (N1) in medial occipital cortex and a positive potential (P1) over bilateral occipital cortex. The second time window implicated a positive distribution in medial occipital cortex (P2) from 188 ms and 238 ms. The late time window from 254 ms to 386 ms with a negative potential (N3) in frontal and parietal cortex corresponded to a higher-order cognitive processing (Figure 3B). Meanwhile, in light of GMD results, topographies among six dimotif lattice stimuli showed a similar pattern regarding the three stages of differentiating six AR conditions by proximity and similarity principles (86-130 ms, 188-238 ms, and 254-386 ms; Figure 3C).

**Figure 3.**
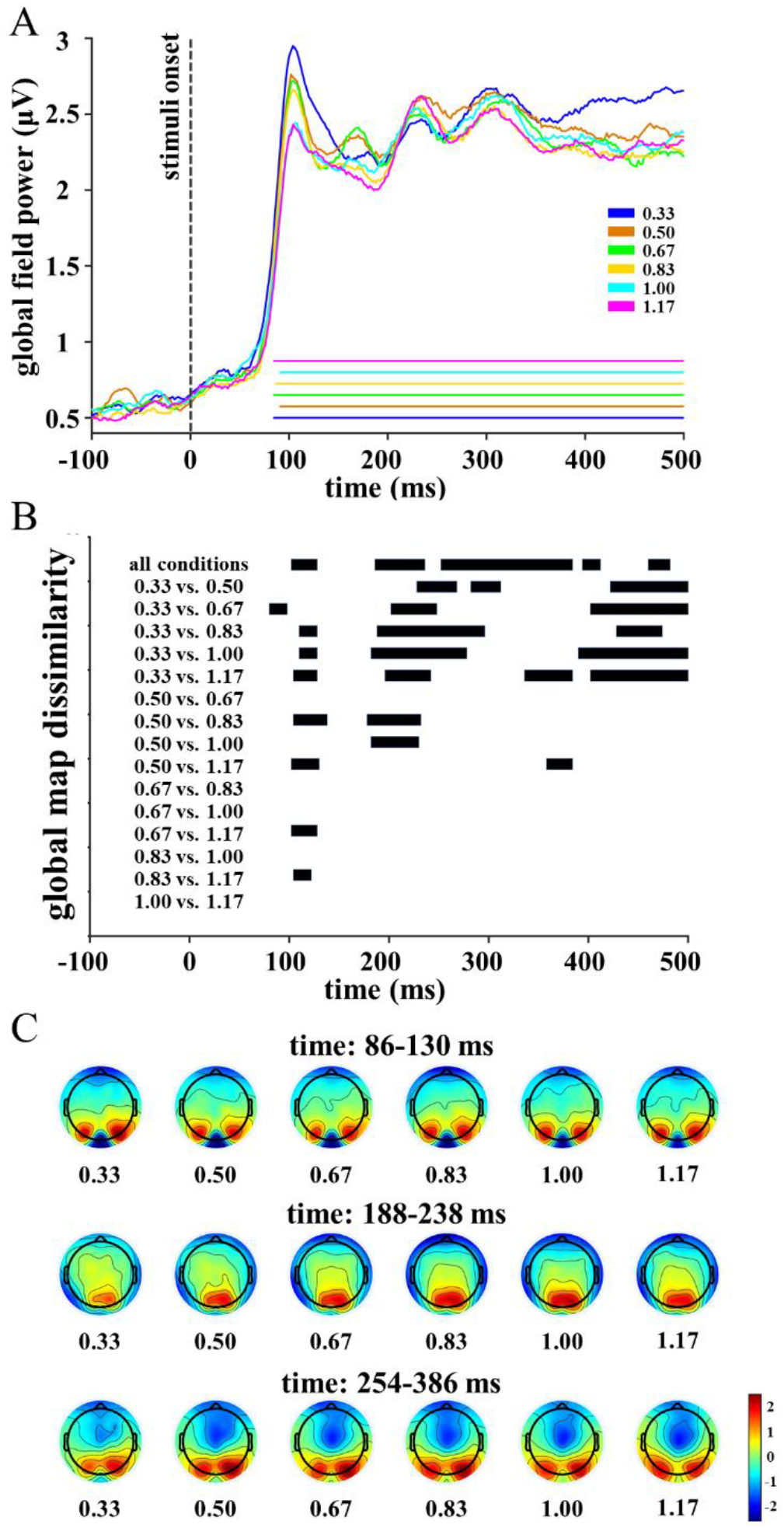
Event-related responses to different ARs exhibited differences in electric field strength and topographic distribution. (A) GFP for each AR pattern (averaged across participants). The time segments whose GFP significantly differed from pre-stimulus baseline were marked with colored straight lines for each AR pattern. (B) The time segments of significant GMD across six-AR patterns were indicated in black bars. The top row depicts the significant main effect of ARs, while the following rows represented all significant pairwise comparison results across six ARs. (C) Topography among six-AR patterns in three time windows (86-130 ms, 188-238 ms, and 254-386 ms). For each topography, EEG signals were averaged across time points within a specific time window. Color bar denotes the voltage value (μV).

### Source estimation results

To identify the brain representational distances and activation patterns associated with proximity and similarity grouping preferences, we calculated the multi-dimensional scaling and the cortical generators of the stimulus-evoked responses for dimotif lattice patterns at three time windows (86-130 ms, 188-238 ms, and 254-386 ms). The results of multidimensional scaling indicated that the initial representation of dimotif lattice patterns in participants’ brain was arranged according to the AR values (86-130 ms). Then, based on the visual processing of elements’ features and spatial locations, the representational distances of dimotif lattice patterns were rearranged. Especially, the ARs of 0.67 and 0.83 were transformed to those close to the representation of similarity grouping preference (i.e., AR=1.00/1.17) (188-238 ms). Finally, at the final stage of global perceived orientation decision-making (254-386 ms), the representational distances showed comparable patterns to behavioral responses (Figure 4A). In addition, brain areas were found associated with the dissociative processing of grouping preferences. First, the visual areas in cuneus and middle occipital gyrus [Brodmann areas (BA) 17, 18, and 19] were engaged in the early processing of spatial relationships between local elements with a time window of 86-130 ms. The attentional post-perceptual operations employed the inferior occipital gyrus and fusiform gyrus [BA 17, 18, 19], indexing the discrimination and selection of stimuli features which were required for completing shape similarity identification from 188 to 238 ms. At the third stage (254-386 ms), the postcentral gyrus (BA 7) was linked to the discrimination in confidence for perceptual grouping decisions (Figure 4B and Table 1).

**Table 1.**
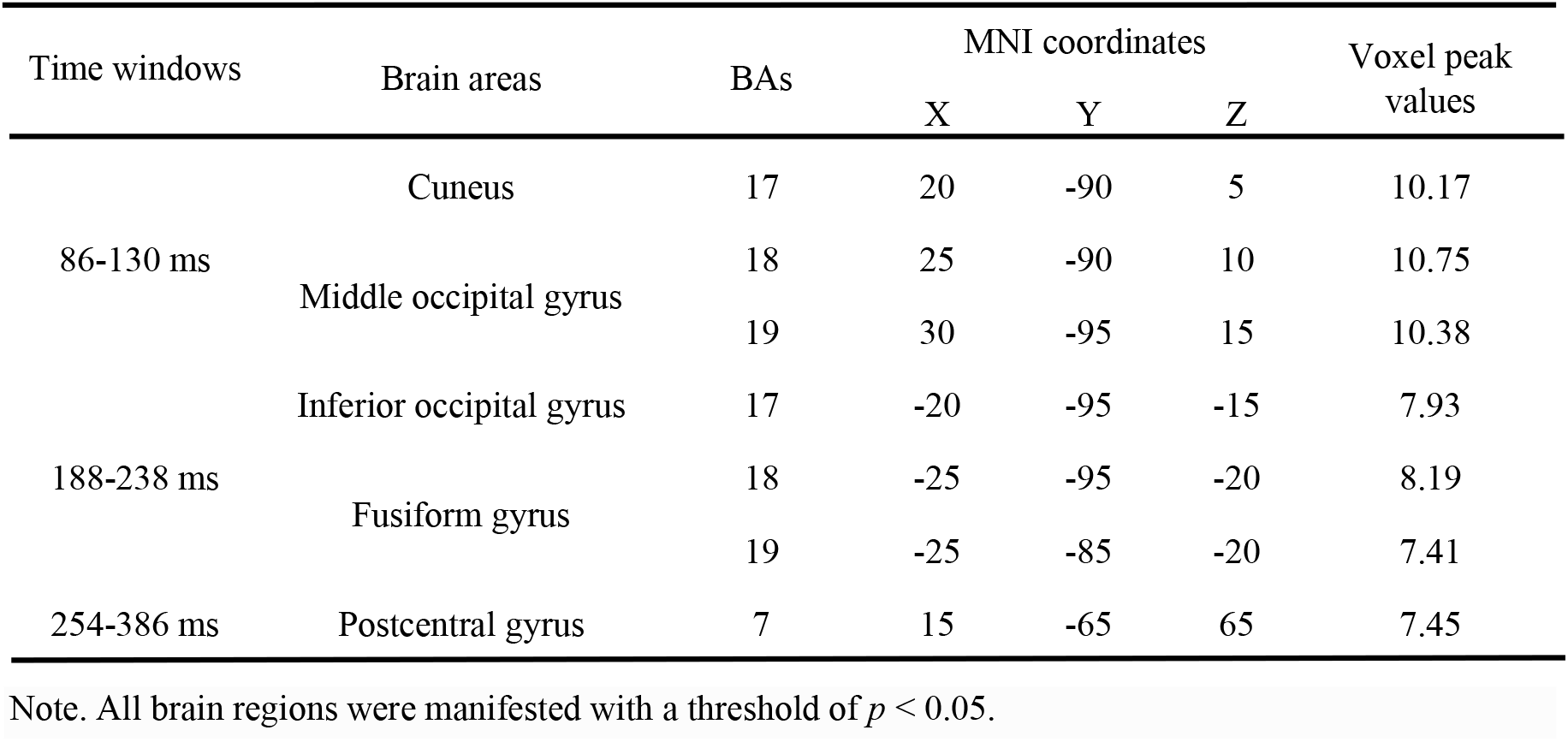
Brain regions showing significant differences across three time windows between max proximity vs. max similarity grouping preference.

**Figure 4.**
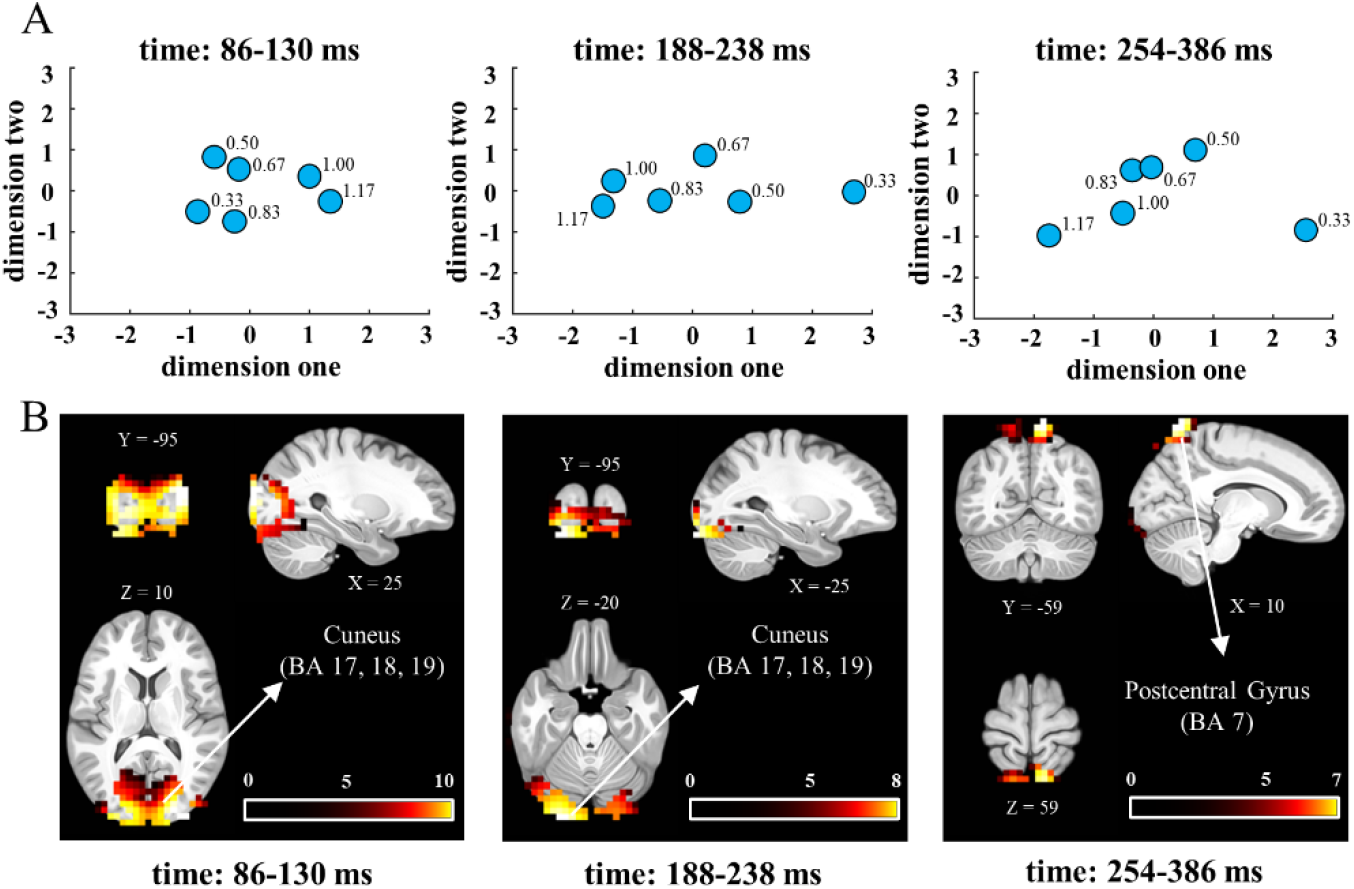
The spatial layout of six ARs and EEG source estimation of proximity and similarity across three time windows. (A) Multi-dimensional scaling provides a visual representation that projects the distances among six AR conditions to examine the representational distances. (B) Estimations of the neural sources in the dissociative processing of two grouping principles underlying the stimulus-evoked responses, including the cuneus and middle occipital gyrus (BA 17, 18, and 19; 86-130 ms), the inferior occipital gyrus and fusiform gyrus (BA 17, 18 and 19; 188-238 ms), and the postcentral gyrus (BA 7; 254-386 ms) (*p* < 0.05, single *t*-test).

### Multi-class decoding and representational similarity results

While the global electric field analysis failed to sufficiently accommodate the discrimination in each dimotif lattice pattern, the dimotif lattice pattern might contain mixed proximity and similarity grouping preferences per se. Alternatively, time-resolved MVPA was employed to evaluate whether the single-trial, instantaneous topographical pattern of EEG activity carries the information about proximity and similarity grouping preference. In the multi-class analysis, grouping preference in proximity and similarity principle was also decoded at three time windows after stimulus onset at group level, which were significant above chance level: 98-146 ms (*t*_(25)_ = 4.202, *p* < 0.001, 95% CI [0.996, 2.912]), 164-190 ms (*t*_(25)_ = 4.230, *p* < 0.001, 95% CI [0.763, 2.211]), and 248-500 ms (*t*_(25)_ = 6.710, *p* < 0.001, 95% CI [1.354, 2.553]). Specifically, those statistics were generated from averaged classification accuracies between normal label and random label during the respective time windows and following-up paired *t*-test comparisons. The results suggested a three-stage process of early, middle, and late perceptual grouping in the human brain, which therefore provided sufficient and accurate time-resolved information for discriminating between grouping preferences in proximity and similarity (Figure 5B).

**Figure 5.**
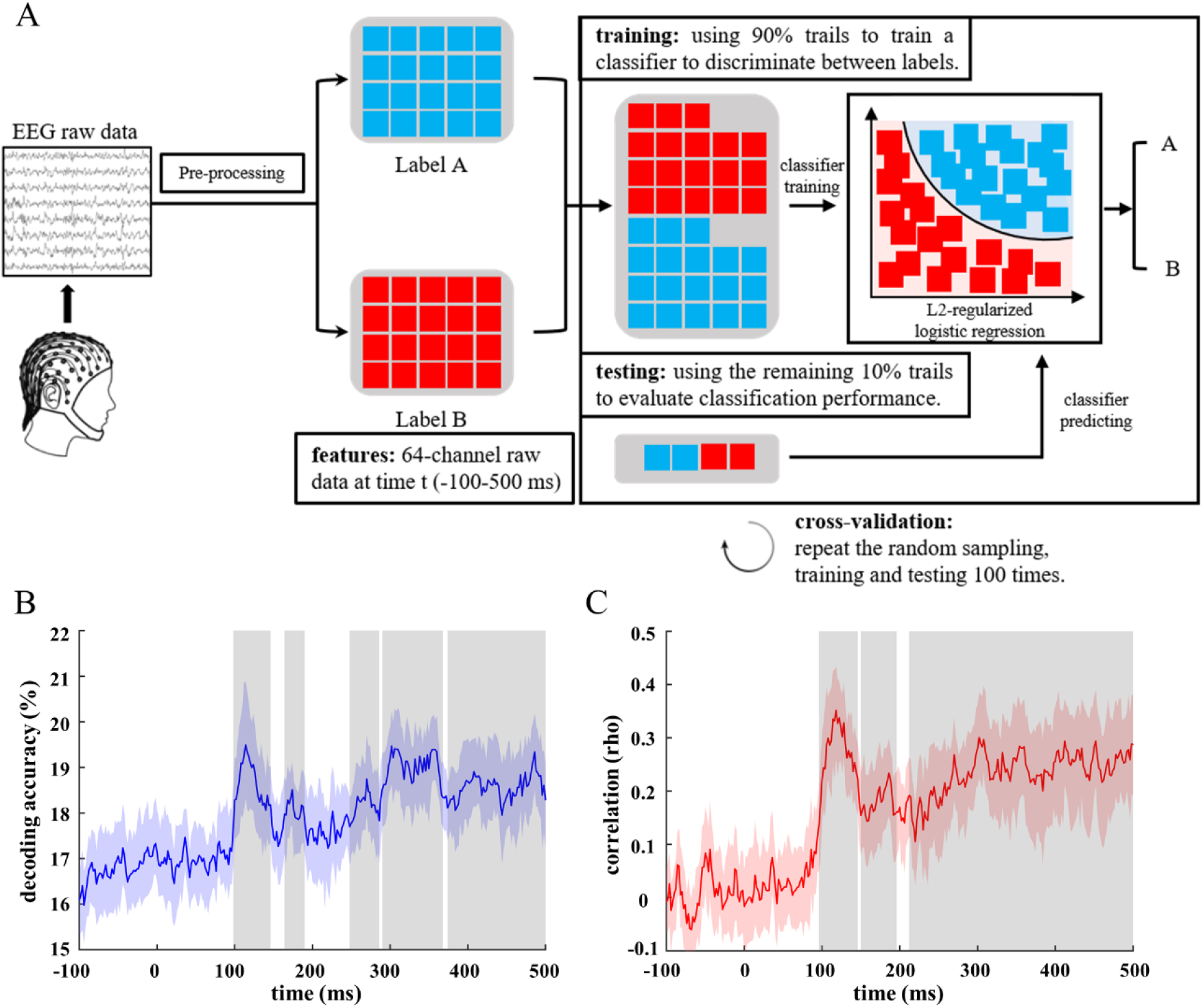
The procedure of MVPA and the visualization of multi-class decoding and representational similarity analysis results. (A) For a given participant and time point, a random sample of 90% of the trails was used to train a classifier to discriminate between brain responses (EEG scalp topography) associated with two (one-versus-one classification) or six (multi-class decoding) ARs. Classification performance was then evaluated by using the remaining 10% of the trails. The entire procedure was repeated for 100 cross-validations. (B) Multi-class decoding: classifiers were trained at each time point among six ARs to reveal when ARs could be dissociated from each other. (C) Representational similarity analysis: behavioral representational similarity matrices were correlated with neural representational similarity matrices at each time point to estimate the similarity between AR representations in perceptual space and neural space. Colored curves denote decoding accuracy across participants [colored shaded area: bootstrapped 95% confidence interval; light gray shaded area: the period of significant decoding at the group level (*p*s < 0.05 lasting for > 20 ms)].

To further quantify the resemblance between behavioral performance and brain responses, we explored the representational similarity between subjective and neural perceptual grouping representations. Since the multi-class decoding results were consistent with behavioral patterns, the dynamic association between participants’ perceptual space and neural space could be obtained. Correlational analyses revealed that behavioral and neural integration were significantly correlated at three time windows: 96-146 ms (*t*_(25)_ = 8.734, *p* < 0.001, 95% CI [0.209, 0.338]), 150-198 ms (*t*_(25)_ = 5.057, *p* < 0.001, 95% CI [0.109, 0.259]) and 212-500 ms (*t*_(25)_ = 6.458, *p* < 0.001, 95% CI [0.159, 0.309]) (Figure 5C). This pattern was comparable to the three stages at which the neuronal signal carried information about grouping preference as indicated by the multi-class analysis (Figure 5B). This line of findings confirmed that the information used by the decoding classifiers formed the basis of perceptual grouping decision-making for each participant.

### One-vs-one decoding results

To further examine how the global perceived orientation varies across different stimuli patterns, we performed one-vs-one classification for all paired dimotif lattice patterns. As shown in Figure 6, pairwise classifications yielded significant accuracies above chance level at certain time points across the trials (*p*s < 0.05 lasting for >20 ms). Pronounced decoding performance between the AR of 0.33 and 1.00/1.17, roughly from 90 to 500 ms, indicated a significant discrimination between stronger proximity and stronger similarity grouping effect (Figure 6, blue curves). Meanwhile, the significant decoding performance between 0.33 AR and 0.50/0.67/0.83 was found at the late time window (322-344 ms for decoding 0.33 vs. 0.50; 294-340 ms, 344-372 ms, and 408-430 ms for decoding 0.33 vs. 0.67; 228-282 ms and 286-500 ms for decoding 0.33 vs. 0.83), suggesting that the different effects between stronger proximity and weaker proximity or between stronger proximity and weaker similarity grouping influenced perceptual grouping decision (Figure 6, blue curves). Interestingly, besides the late time window, weaker proximity grouping effect (AR = 0.50) was distinguished from stronger similarity grouping effect (AR = 1.00/1.17) at an early time window (104-142 ms for decoding 0.50 vs. 1.00; 96-144 ms for decoding 0.50 vs. 1.17; Figure 6, pink curves). Furthermore, significant effect was only found at the early time window between stronger similarity (AR = 1.17) and weaker similarity grouping (AR = 0.67, 128-148 ms, green curve; AR = 0.83, 98-118 ms, yellow curve). Yet, no significant effect was detected between the adjacent AR values of similarity grouping preference (*p*s > 0.05, red curve).

**Figure 6.**
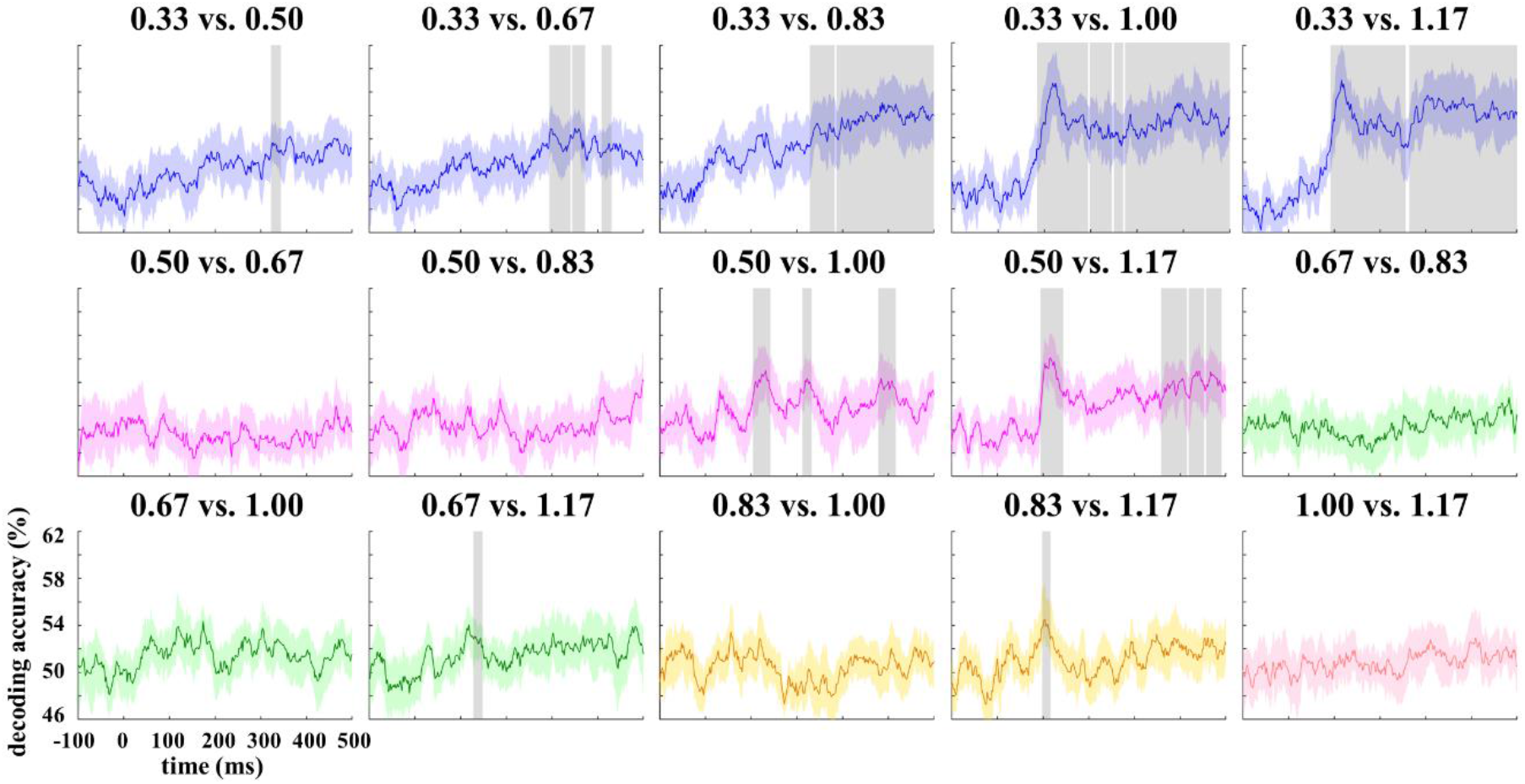
One-vs-one decoding reveals that different ARs were represented by distinctive neural patterns. For one-vs-one decoding, classifiers were trained at each time point to provide which time points were different between paired ARs. Colored curves indicate decoding accuracy averaged across participants [colored shaded area: bootstrapped 95% confidence interval; light gray shaded area: the period of significant decoding at the group level (*ps* < 0.05 lasting for > 20 ms)]. Various colors were employed to denote differing comparisons.

### Temporal generation matrix results

Furthermore, the neural template analysis confirmed the differing effects between stronger proximity and stronger similarity grouping, and meanwhile the overlapping patterns in the dimotif lattice pattern between weaker proximity and weaker similarity grouping co-process. In particular, when paired classification was conducted between stronger proximity (AR = 0.33) and stronger similarity (AR = 1.00/1.17, Figure 7), unified and larger cluster could be identified within the three time windows. This could suggest that decoding performance for the sub-processes reflected certain common neural mechanisms. However, the large cluster was only detected at a late time window when paired classification was conducted between stronger proximity (AR = 0.33) and weaker similarity (AR = 0.50), or between stronger proximity (AR = 0.33) and weaker similarity grouping preference (AR = 0.67/0.83). In addition, significant clusters were found at both early and late time windows between stronger similarity (AR = 1.00/1.17) and weaker proximity grouping (AR = 0.50). Yet, the small clusters were only found at the late time window between weaker similarity (AR = 0.67) and the other similarity grouping preference (AR = 0.83/1.00/1.17). No significant cluster was found among similarity grouping except the AR value of 0.67. Thus, the neural patterns carrying the information for the persistent decoding of grouping preference in proximity or similarity were altering from one moment to the next in the three time windows.

**Figure 7.**
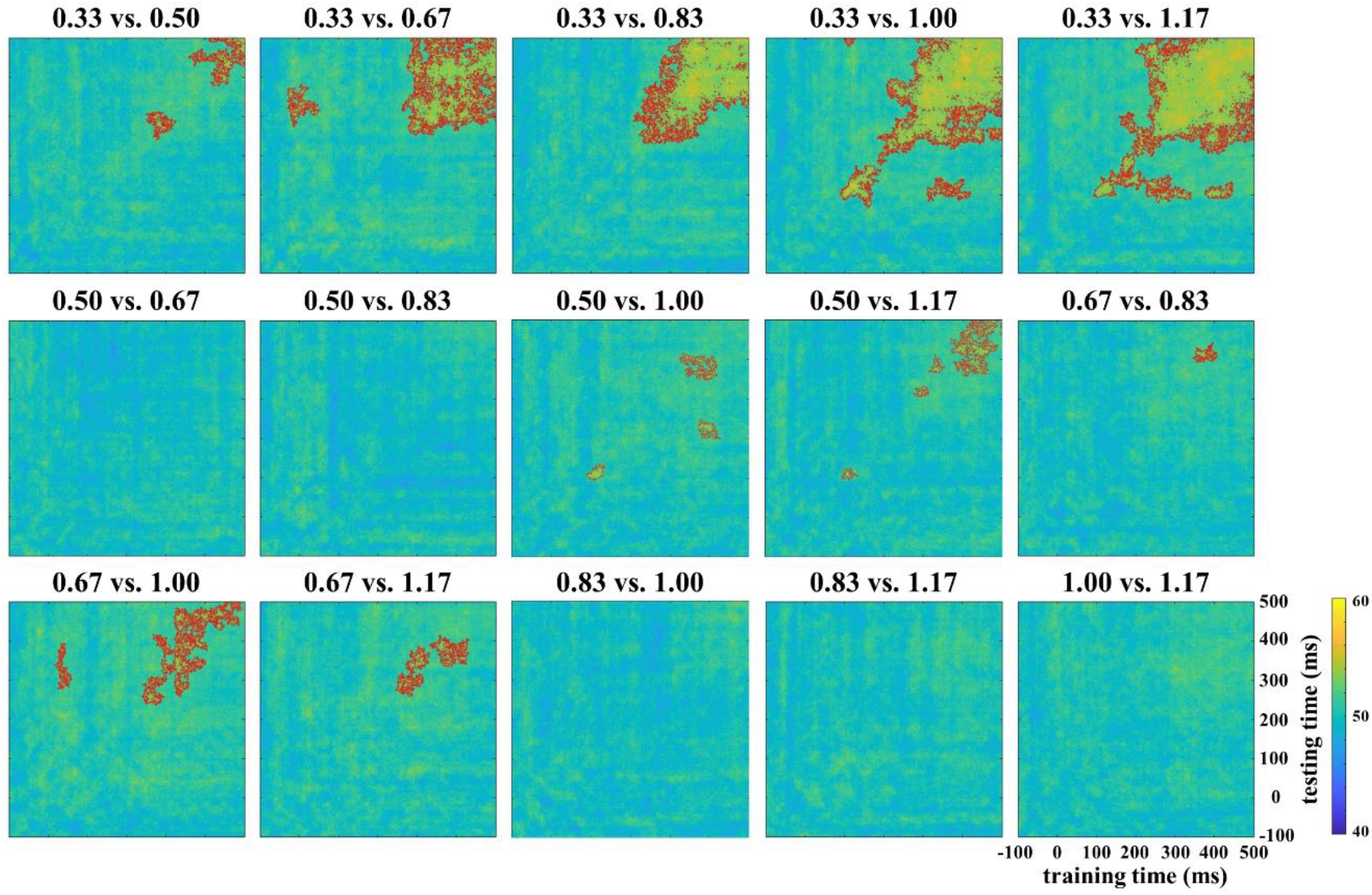
Neural template analysis testing for the generalization of decoding across time and ARs based on one-vs-one classification. Classifiers were first trained to discriminate one AR versus the other at a given time point (training time *t*_*x*_). This classifier was then tested by using it to predicate two ARs at all time points (testing time *t*_*y*_). Significant decoding along the main diagonal indicates that two ARs are represented by similar neural patterns at the same latency. On the contrary, significant decoding along the off-diagonal dimension indicates that two ARs are represented by similar neural patterns yet at different latencies (multiple comparisons: FDR, *p* < 0.05). Color bar denotes the decoding accuracy (%).

## Discussion

A whole picture of the dissociative processing between two perceptual grouping principles in humans was depicted in both perceptual and neural domains in the current study. First, psychophysical results revealed that as AR values increased, more global perceived judgements for similarity orientation were made. This pattern was in line with the findings from <u>Kubovy and Van Den Berg (2008)</u> and <u>Wei et al. (2018)</u>, which showed that similarity grouping effect is enhanced with decreased proximity grouping effect when the two unstable grouping cues were showed simultaneously. Importantly, we quantified the perception gap between proximity and similarity principles which approximates 1.60°. In addition, the precise distance after converting from visual angle space to physical space (1.54 cm) was close to the value from the seminal work of this field, which demonstrated that “… it may be said that Similarity equals about 1.5 cms. of Proximity” (<u>Rush, 1937</u>). This could somehow validate our paradigm, which decreased the risk of data contamination due to participants’ subjective reports and thus obtained relatively reliable results of the relationship between the two grouping principles.

With regard to the global electric field differences among six-stimulus patterns, the time at which perceptual grouping information was available coincided with the latency of the first visual-evoked response at around 90 ms. It suggested that perceptual grouping information could initially be represented by one of the most elementary features. In visual system, the initial event-related response is associated with the representation of visual features and a fast category-selective activity, when the visual system is highly optimized for the processing of natural scenes (<u>Wang et al., 2012</u>; <u>Kaneshiro et al., 2015</u>). Existing EEG studies have identified that the early component (P1) was only associated with proximity-defined grouping (<u>Han, 2004</u>; <u>2005b</u>). In light of the comparable patterns we obtained, our GFP results therefore extended this line of evidence by demonstrating that the human brain might also initially manifest a proximity preference with the presence of the both grouping cues, while the final grouping decision-making is similarity preference. Importantly, GMDs confirmed the hypothesis for three time windows (86-130 ms, 188-238 ms and 254-386 ms), indexing the processing of distinguishing two perceptual grouping principles. The current findings further indicated that different sub-regions within the parieto-occipital and fronto-parietal networks are activated by proximity and similarity preference to different extents across three time windows. Note that these findings were compatible with different proposals for the extensive network, which argued the frontal, parietal and occipital cortex were commonly engaged in the processing of proximity and similarity grouping (<u>Han and Humphreys, 2007</u>; <u>Seymour et al., 2008</u>).

Further, multi-dimensional scaling and source estimation provided a visualization of likely representational distance and predominant sources in the human brain for the three-stage processing of grouping preference dissociation. The discrimination primarily involves the middle occipital cortex at an early stage (86-130 ms), the inferior occipital cortex and fusiform cortex at middle-stage (188-238 ms), and the parietal cortex at late stage (254-386 ms). These areas have been widely recognized to be associated with perceptual grouping processing in previous functional neuroimaging studies. The middle occipital cortex in cuneus, in particular, was linked to the proximity-defined larger-scale organization of discrete visual elements in the visual field (<u>Han et al., 2005a</u>). Our results validated the previous findings by using multi-dimensional scaling analysis, which showed that the pattern distances are arranged consistently with the strength of proximity effect in the early stage. Then, in the middle stage, a stronger grouping-related activation was observed in the inferior occipital and fusiform gyrus (BA 17, 18 and 19). This pattern was consistent with fMRI studies which specified the functional contribution of object recognition (<u>Joseph, 2001</u>; <u>Sim et al., 2015</u>). Furthermore, multi-dimensional scaling showed that the aspect ratios (AR = 0.67 and 0.83) were closer to the similarity set, which were initially closer to the proximity set. This might reflect a feedback mechanism from the lateral occipital cortex, the inferior occipital, and fusiform gyrus, to V1/V2 selecting the object features and processing the object recognition (<u>Chen et al., 2021</u>). Finally, the parietal cortex is considered as the area for spatial attention (<u>Kim et al., 2017</u>; <u>Grassi et al., 2018</u>) and more engagements were found during a feature integration task (<u>Shafritz et al., 2002</u>; <u>Freedman and Ibos, 2018</u>). The current findings in late component therefore implicated a further distinction between two grouping information and final decision-making based on spatial attention and feature integration.

To identify the subtle temporal representation of the dissociative processing across three time windows, we trained classifiers on the data collected from different grouping preference conditions, respectively. The classifiers were then tested by looking at the decoding performance and neural template across all electrodes for each time point. Interestingly, the neural response patterns obtained from multi-class MVPA of EEG recordings corroborated results of global electric field. With an early onset and relatively short peak, the classification performance suggested that earlier sources of dissociative information came from the dynamic visual representations (<u>Guo et al., 2019</u>). For each grouping preference pattern, significant decoding performance between differing preferences could cover more than one time segment. Interestingly, paired comparisons between any two consecutive AR values were pronounced in the early time window for the pairs of similarity grouping preference, while prominent in the late time window for the proximity pairs. The pattern manifested in the early time window could corroborate the notion that the distinct processing between two grouping principles initially employed proximity to group the ordered and discrete visual elements. Meanwhile, this line of finding reflected the proximity-defined large-scale organization (<u>Han et al., 2005a</u>; <u>2005b</u>), even in the paired conditions of similarity grouping preference. Importantly, the late time window (>260 ms) reported in previous EEG studies also elaborated the higher-order cognitive function integrating perception to action (<u>Deslandes et al., 2005</u>; <u>Takacs et al., 2020</u>). The current findings further reveled that more cognitive resources were engaged to discriminate differing stimuli and determine global perceived orientation when distant AR values were compared.

Furthermore, in light of the correlational results, perceptual and neural space converged after first stimulus-evoked response (96-144 ms), demonstrating that the distinguished perceptual grouping information encoded in MVPA laid the basis for perceptual grouping decision-making. The more similarly any two dimotif lattice patterns were represented in MVPA, the more confused the participants might feel in the process of perceptual grouping decision-making. To our knowledge, our study is the first to show that the subjective discriminability of perceptual grouping principles is related to their neural dissimilarity, thus manifesting a robust neural-perceptual mapping.

In summary, the present study was among the first to apply MVPA to decode the processing of two perceptual grouping information, particularly with an emphasis on the representational similarity of perceptual space and neural space. Notably, the psychophysical responses reported here were in strong alignment with previous behavioral studies and we further quantified the perception gap between proximity and similarity principles. Our decoding results showed that the human visual system dissociating two perceptual grouping information is encoded at three stages of perceptual grouping. The grouping starts with the proximity-defined arrangement of local discrete visual elements in the middle occipital cortex, followed by the feature selection and spatial attention modulating two perceptual grouping cues in the inferior occipital cortex and fusiform cortex. At last, it ended with the higher cognitive integration for spatial information and feature conjunction to determine the decision-making in the parietal cortex. Thus, our findings offer fundamental insights into the mental separation of two perceptual grouping processing and its relation to ultimate perceptual grouping-related decisions. We anticipate that our methods and results will ignite further investigations using time-resolved whole-brain responses to understand other perceptual grouping principles, which could also contribute to other sensory perceptual grouping processing.

## References

Carlson T, Tovar DA, Alink A, Kriegeskorte N (2013) Representational dynamics of object vision: the first 1000 ms. Journal of vision 13:1–1.

Carther-Krone TA, Lawrence-Dewar JM, Shomstein S, Nah JC, Collegio AJ, Marotta JJ (2020) Neural Correlates of Perceptual Grouping Under Conditions of Inattention and Divided Attention. Perception 49:495–514.

Cauchoix M, Crouzet SM, Fize D, Serre T (2016) Fast ventral stream neural activity enables rapid visual categorization. NeuroImage 125:280–290.

Chen S, Weidner R, Zeng H, Fink GR, Müller HJ, Conci M (2021) Feedback from lateral occipital cortex to V1/V2 triggers object completion: Evidence from functional magnetic resonance imaging and dynamic causal modeling. Human Brain Mapping.

Cichy RM, Pantazis D (2017) Multivariate pattern analysis of MEG and EEG: A comparison of representational structure in time and space. NeuroImage 158:441–454.

Cichy RM, Pantazis D, Oliva A (2014) Resolving human object recognition in space and time. Nature neuroscience 17:455–462.

Crouzet SM, Busch NA, Ohla K (2015) Taste quality decoding parallels taste sensations. Current Biology 25:890–896.

Delorme A, Makeig S (2004) EEGLAB-Open Source Matlab Toolbox for Electrophysiological Research. Journal of Neuroscience Methods 134:9–21.

Deslandes A, Veiga H, Cagy M, Piedade R, Pompeu F, Ribeiro P (2005) Effects of caffeine on the electrophysiological, cognitive and motor responses of the central nervous system. Brazilian Journal of Medical and Biological Research 38:1077–1086.

Fan R-E, Chang K-W, Hsieh C-J, Wang X-R, Lin C-J (2008) LIBLINEAR: A library for large linear classification. the Journal of machine Learning research 9:1871–1874.

Fang F, Kersten D, Murray SO (2008) Perceptual grouping and inverse fMRI activity patterns in human visual cortex. Journal of vision 8:2–2.

Freedman DJ, Ibos G (2018) An integrative framework for sensory, motor, and cognitive functions of the posterior parietal cortex. Neuron 97:1219–1234.

Grassi PR, Zaretskaya N, Bartels A (2018) A generic mechanism for perceptual organization in the parietal cortex. Journal of Neuroscience 38:7158–7169.

Grünbaum B, Shephard GC (1987) Tilings and patterns: Courier Dover Publications.

Guo LL, Nestor A, Nemrodov D, Frost A, Niemeier M (2019) Multivariate Analysis of Electrophysiological Signals Reveals the Temporal Properties of Visuomotor Computations for Precision Grips. Journal of Neuroscience 39:9585–9597.

Han S (2004) Interactions between proximity and similarity grouping: An event-related brain potential study in humans. Neuroscience Letters 367:40–43.

Han S, Humphreys GW (2007) The fronto-parietal network and top-down modulation of perceptual grouping. Neurocase 13:278–289.

Han S, Jiang Y, Mao L, Humphreys GW, Gu H (2005a) Attentional modulation of perceptual grouping in human visual cortex: Functional MRI studies. Human Brain Mapping 25:424–432.

Han S, Jiang Y, Mao L, Humphreys GW, Qin J (2005b) Attentional modulation of perceptual grouping in human visual cortex: ERP studies. Human Brain Mapping 26:199–209.

Joseph JE (2001) Functional neuroimaging studies of category specificity in object recognition: a critical review and meta-analysis. Cognitive, Affective, & Behavioral Neuroscience 1:119–136.

Kaneshiro B, Perreau Guimaraes M, Kim H-S, Norcia AM, Suppes P (2015) A representational similarity analysis of the dynamics of object processing using single-trial EEG classification. Plos one 10:e0135697.

Kim Y-J, Tsai JJ, Ojemann J, Verghese P (2017) Attention to multiple objects facilitates their integration in prefrontal and parietal cortex. Journal of Neuroscience 37:4942–4953.

Kriegeskorte N, Mur M, Bandettini PA (2008) Representational similarity analysis-connecting the branches of systems neuroscience. Frontiers in systems neuroscience 2:4.

Kubovy M, Van Den Berg M (2008) The whole is equal to the sum of its parts: A probabilistic model of grouping by proximity and similarity in regular patterns. Psychological review 115:131.

Kurylo DD, Waxman R, Kidron R, Silverstein SM (2017) Visual training improves perceptual grouping based on basic stimulus features. Attention, Perception, & Psychophysics 79:2098–2107.

Luna D, Montoro PR (2011) Interactions between intrinsic principles of similarity and proximity and extrinsic principle of common region in visual perception. Perception 40:1467–1477.

Luna D, Villalba-García C, Montoro PR, Hinojosa JA (2016) Dominance dynamics of competition between intrinsic and extrinsic grouping cues. Acta psychologica 170:146–154.

Merigan WH, Nealey TA, Maunsell J (1993) Visual effects of lesions of cortical area V2 in macaques. Journal of Neuroscience 13:3180–3191.

Murray MM, Brunet D, Michel CM (2008) Topographic ERP analyses: a step-by-step tutorial review. Brain topography 20:249–264.

Murray SO, Schrater P, Kersten D (2004) Perceptual grouping and the interactions between visual cortical areas. Neural Networks 17:695–705.

Nikolaev AR, Gepshtein S, Kubovy M, Van Leeuwen C (2008) Dissociation of early evoked cortical activity in perceptual grouping. Experimental brain research 186:107–122.

Pascual-Marqui RD (2002) Standardized low-resolution brain electromagnetic tomography (sLORETA): technical details. Methods Find Exp Clin Pharmacol 24:5–12.

Pauli WM, Nili AN, Tyszka JM (2018) A high-resolution probabilistic in vivo atlas of human subcortical brain nuclei. Scientific data 5:1–13.

Quinlan PT, Wilton RN (1998) Grouping by proximity or similarity? Competition between the Gestalt principles in vision. Perception 27:417–430.

Rush GP (1937) Visual grouping in relation to age. Archives of Psychology (Columbia University).

Schütt HH, Harmeling S, Macke JH, Wichmann FA (2016) Painfree and accurate Bayesian estimation of psychometric functions for (potentially) overdispersed data. Vision research 122:105–123.

Seymour K, Karnath H-O, Himmelbach M (2008) Perceptual grouping in the human brain: common processing of different cues. Neuroreport 19:1769–1772.

Shafritz KM, Gore JC, Marois R (2002) The role of the parietal cortex in visual feature binding. Proceedings of the National Academy of Sciences 99:10917–10922.

Sim E-J, Helbig HB, Graf M, Kiefer M (2015) When action observation facilitates visual perception: activation in visuo-motor areas contributes to object recognition. Cerebral cortex 25:2907–2918.

Stoll S, Finlayson N, Schwarzkopf DS (2017) The topographic representation of global object perception in human visual cortex. Journal of Vision 17:747–747.

Takacs A, Zink N, Wolff N, Münchau A, Mückschel M, Beste C (2020) Connecting EEG signal decomposition and response selection processes using the theory of event coding framework. Human Brain Mapping 41:2862–2877.

Villalba-García C, Santaniello G, Luna D, Montoro P, Hinojosa J (2018) Temporal brain dynamics of the competition between proximity and shape similarity grouping cues in vision. Neuropsychologia 121:88–97.

Wagemans J, Elder JH, Kubovy M, Palmer SE, Peterson MA, Singh M, von der Heydt R (2012) A century of Gestalt psychology in visual perception: I. Perceptual grouping and figure–ground organization. Psychological bulletin 138:1172.

Wang C, Xiong S, Hu X, Yao L, Zhang J (2012) Combining features from ERP components in single-trial EEG for discriminating four-category visual objects. Journal of neural engineering 9:056013.

Wannig A, Stanisor L, Roelfsema PR (2011) Automatic spread of attentional response modulation along Gestalt criteria in primary visual cortex. Nature neuroscience 14:1243–1244.

Wei N, Zhou T, Chen L (2018) Objective measurement of gestalts: Quantifying grouping effect by tilt aftereffect. Behavior research methods 50:963–971.

Wertheimer M (1922) Untersuchungen zur Lehre von der Gestalt. Psychologische forschung 1:47–58.

Wertheimer M (1923) Untersuchungen zur Lehre von der Gestalt II [Investigations on the Gestalt Theory II]. Psycologische Forschung, 4. In.

